# Metanalysis of genome-wide association studies for panic disorder suggest pathways and mechanisms of pathogenesis

**DOI:** 10.1101/326017

**Authors:** Rhayra Xavier do Carmo Silva, Sueslene Prado Rocha, Dainara Pereira dos Santos Souza, Monica Gomes Lima-Maximino, Caio Maximino

## Abstract

Panic disorder (PD) is characterized by abrupt surges of intense fear and distress. There is evidence for a genetic component in this disorder. We ran a meta-analysis of genome-wide association studies of patients with PD, and found 25 single-nucleotide polymorphisms that were associated with the disorder. Causal gene prediction based on these polymorphisms uncovered 20 hits. Exploratory analyses suggested that these genes formed interactor networks, which was enriched in signaling pathways associated with immune and inflammatory responses, as well as growth factors and other developmental mediators. A subset of genes is enriched in limbic regions of the human brain and in microglia and myelinating oligodendrocytes of mice. While these genes were not associated with relevant neurobehavioral phenotypes in mutant mice, expression levels of several causal genes in the amygdala, prefrontal cortex, hippocampus, hypothalamus, and adrenal gland of recombinant mouse strains was associated with endophenotypes of fear conditioning. Drug repositioning prediction was unsuccessful, but this does not discard these genes and pathways as targets for investigational drugs. In general, *ASB3*, *EIF2S2, RASGRF2*, and *TRMT2B* (and its coded proteins) emerged as interesting targets for mechanistic research on PD. These exploratory findings point towards hypotheses of pathogenesis and neuropharmacology that need to be further investigated.

## 1 Introduction

Panic disorder (PD) is characterized by repeated and unpredictable panic attacks, usually associated with the development of anticipatory anxiety and avoidance strategies that can culminate in agoraphobia (American Psychiatric Association, 2013). Patients with PD have high rates of medically unexplained symptoms that produce a burden on healthcare services (Coley et al., 2009). While highly vulnerable to environmental and life-event causes, panic disorder have a significant genetic component. There is evidence of familial aggregation and moderate heritability of PD (~44%)(Merikangas and Pine, 2002). Many genetic studies have tried to identify linkage or association of specific anxiety disorders with genomic regions or specific genes, but the success rate of such studies is low (Gratacòs et al., 2007). Different approaches to understand the genetic vulnerabilities associated with anxiety disorders have been proposed, from quantitative trait loci studies and gene expression profiling in animal models (Sokolowska and Hovatta, 2013), knockout studies in mice (Wood and Toth, 2001), and genome-wide association studies (GWAS) in humans (Binder, 2012). Combining different sources of genetic information may help to identify relevant candidate genes and novel drug targets for the treatment of anxiety and stress disorders (Binder, 2012; Hettema et al., 2011; Lotan et al., 2014; Ofrat and Krueger, 2012; Stewart et al., 2015).

An important addition to that approach is the use of computational-informatic approaches to find patterns in expansive data sets (Lotan et al., 2014; Stewart et al., 2015). Data mining approaches are exploratory techniques which can help produce *a posteriori* hypothesis for datasets with a high amount of information. These techniques have been applied to GWAS analyses of psychiatric disorders to not only make sense of the huge amount of data, but also to inform future research on the pathophysiology and pharmacology of these diseases. For example, So and colleagues (So, 2017; So et al., 2016, 2017) used gene-set analyses to propose drug repositioning for depression and anxiety disorders, with good predictive value. Lotan et al. (2014) analyzed single nucleotide polymorphisms (SNPs) derived from GWAS for attention-deficit/hyperactivity disorder, anxiety disorders, autism spectrum disorder, bipolar disorder, major depressive disorder, and schizophrenia, and found that 22% the genes associated with those SNPs are shared between at least two of those groups; they also found that the shared genes are enriched in the postsynaptic density, expressed in immune tissues, and co-expressed in the developing human brain. Wray et al. (2018) found that genes associated with increased risk for major depressive disorder in GWAS are enriched in the frontal and cingulate cortices, as well as in the putamen and nucleus accumbens, of humans, and that these genes were highly expressed in neurons. Using Gene Ontology, they also observed that major depressive disorder-related genes were significantly associated with synaptic genes, genes involved in neuronal morphogenesis, differentiation, and axonogenesis, and genes encoding voltage-gated calcium channels or involved in cytokine and immune response. These examples underline the power of exploratory research using computational approaches to “make sense” of the expansive datasets provided by GWAS studies, and to use these datasets to predict pathophysiology and pharmacology for psychiatric disorders. In the present study, we use a metanalysis of GWAS for PD-related SNPs to derive causal gene predictions. Causal genes were then analyzed in terms of protein-protein interactions, relationship to expression in regions of the adult human brain, cell-specific expression in rodents brain cells, and relationship to neurobehavioral traits in recombinant mice strains to prioritize genes for translational and reverse translational research, as well as to produce pathophysiological insights for PD. We also attempt to predict pharmacological targets by analyzing signaling pathway enrichment and drug repositioning techniques by enrichment analysis.

## 2 Methods

### 2.1 Meta-analysis of case-control association studies and causal gene prediction

Single-nucleotide polymorphisms associated with PD in human patients were mined from a systematic review of the literature using the PubMed search engine. The search strategy was (‘panic disorder’) AND (‘genome-wide association study’). Results were limited to English language papers. The reference lists of the retrieved articles were also reviewed to identify publications on the same topic. The published SNPs within 10 kb of a genomic feature (i.e., gene/transcript biotypes) were selected, and protein-coding features were used for the meta-analysis. Table 1 presents the selected studies. Data were extracted independently and in duplicate by two reviewers (R.X.C.S and S.P.R.) using a standardized data extraction form. Any disagreement was adjudicated by a third author (C.M.).

**Table 1.**
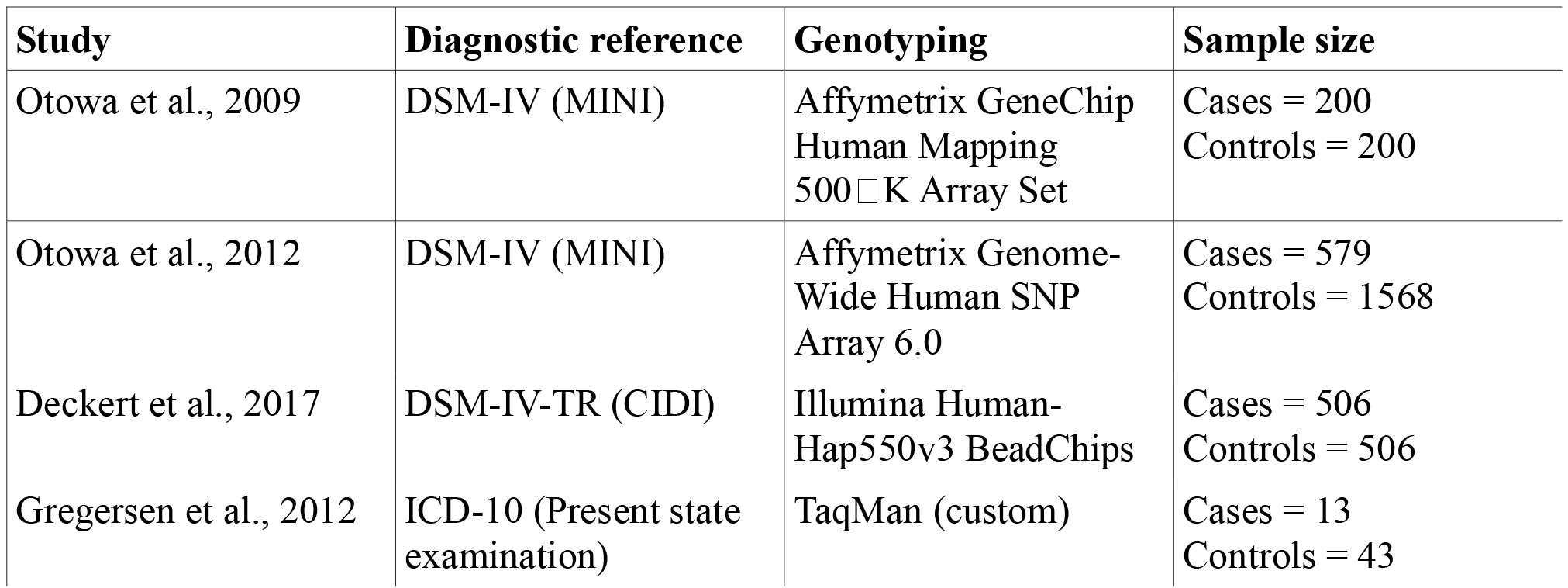
Summary of the studies retrieved for the meta-analysis.

**Table 2.**
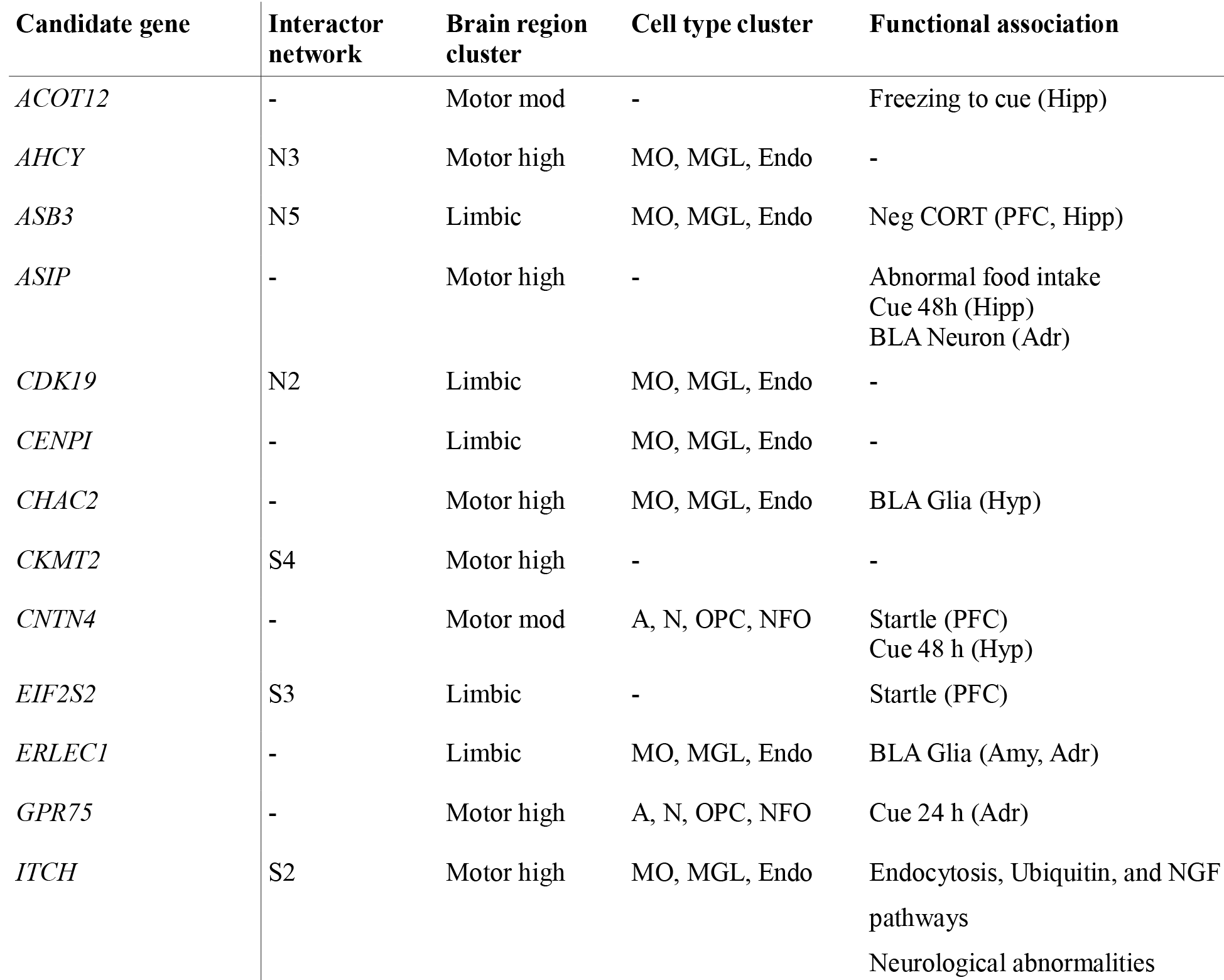
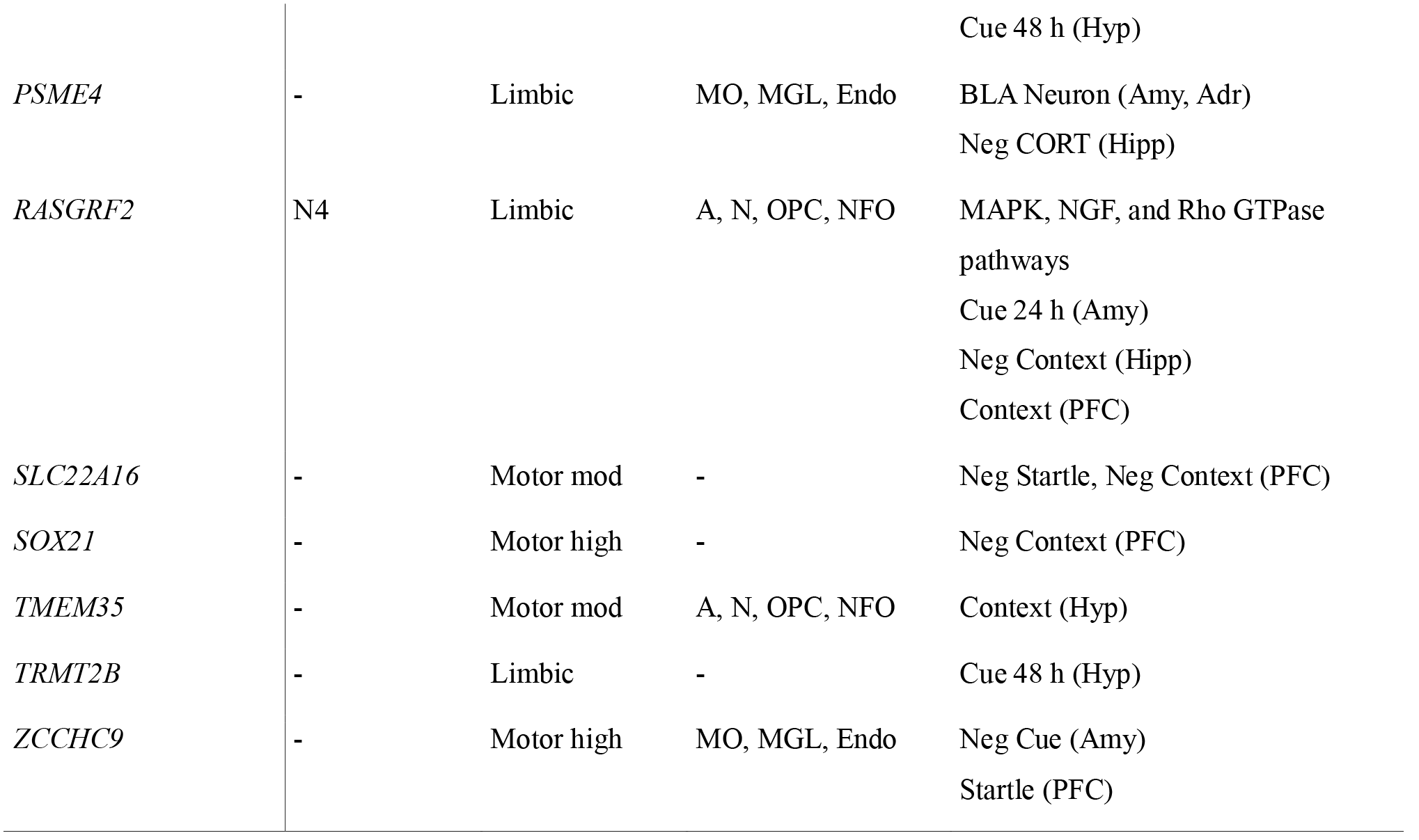
Summary of exploratory findings for the candidate genes found in the meta-analysis and causal gene analysis.

Meta-analyses were conducted on the basis of the fixed-effect model as implemented by METAL by combining summary statistics across data sets (Willer et al., 2010). Since meta-analysis of summary statistics is as powerful as pooling individual-level data across studies (Lin and Zeng, 2009), summarized data (SNP identification, reference allele, non-reference allele, and associated p-values) was used. At the meta-analysis level, summary statistics were filtered for inclusion after meeting a minimum imputation quality score of 0.3, minor allele frequency of greater than 0.1% across studies, and absolute *β* coefficients less than 5. The resulting p-values were corrected for false discovery rates using the R package fdrtool (https://cran.r-project.org/package=fdrtool; Strimmer, 2008). The resulting FDR-corrected trait-associated SNPs (TASs) were filtered based on q-values following a three-sigma rule, with only those TASs that were associated with PD at a q-value smaller than 0.001 retained for posterior analysis.

The resulting TASs were analyzed using the PrixFixe method for causal gene prediction (Taşan et al., 2014); briefly, the method predicts causal genes based on cofunction networks (using data from shared GO terms, genetic interactions, co-expression, transcription factor binding, protein domains, and protein interactions) to identify functionally related genes. This method gives each putative gene a PF score; higher scores equate stronger confidence of the prediction. TASs were analyzed using 500000 b upstream and downstream padding, and a linkage disequilibrium threshold of *r*^2^ = 0.75, corresponding to ‘tightened’ genomic regions (Taşan et al., 2014). Ranks were obtained using version 0.7 of the PrixFixe package for R/Bioconductor (https://llama.mshri.on.ca/~mtasan/GranPrixFixe/html/), and organized per chromosomal region. Only genes with PF scores greater than 0 were retained. These data are available from our GitHub repository (https://github.com/lanec-unifesspa/gwas-pd/tree/master/metanalysis).

PF-ranked genes were then compared to candidate genes associated with panic disorder in human patients mined from the GWASdb2 database (http://jjwanglab.org/gwasdb; Li et al., 2012). These candidate genes are manually mapped to Disease Ontology database ID (DOID) 594, ‘panic disorder’, and PrixFixe scores were calculated automatically by the database. Genes and PF scores were retrieved, and combined with genes found in our own metanalysis through the use of Venn diagrams. Genes which were shared in both datasets were retained for further analysis.

### 2.2 Protein-protein interaction and signalling pathway analyses

The Interaction analysis was made for both high-throughput and small-scale physical, undirected interactions, using PathwayLinker (http://pathwaylinker.org/; Farkas et al., 2012). BioGrid and STRING “exp” databases were used to analyze high-throughput physical interactions, while HPRD was used for small-scale physical interactions between selected proteins and their first neighbors. The resulting set, which was visualized using Cytoscape Web (version 0.8), contained purely physical interactions (i.e., association in complexes, direct interaction, physical interaction, and biochemical co-localization).

Signaling pathway analyses were made using PathwayLinker (http://pathwaylinker.org/; Farkas et al., 2012), using KEGG, Reactome, and SignaLink as databases. Enrichment was considered significant when associated p-values were < 0.001. Both interaction network and signaling pathway data are available from our GitHub repository (https://github.com/lanec-unifesspa/gwas-pd/tree/master/pathwaylinker).

### 2.3 Expression levels in the human brain

Using the Allen Human Brain Atlas (http://human.brain-map.org/; Hawrylycz et al, 2012), gene expression (microarray) data were downloaded for 20 large neuroanatomical divisions (frontal lobe, insula, cingulate gyrus, hippocampal formation, parahippocampal gyrus, occipital lobe, parietal lobe, temporal lobe, amygdala, basal forebrain, globus pallidus, striatum, claustrum, epithalamus, hypothalamus, subthalamus, dorsal thalamus, ventral thalamus, mesencephalon, cerebellar cortex, cerebellar nuclei, basal part of pons, pontine tegmentum, myelencephalon, white matter, sulci & spaces). Data were derived from the brains of healthy human donors, and represented as z-scores. When more than one array was found, the median was calculated; these data are available from our GitHub repository (https://github.com/lanec-unifesspa/gwas-pd/tree/master/Allen). Data were then analyzed with hierarchical cluster with Euclidean distances and average linkage using Cluster 3.0 (University of Tokyo, Japan). Clustering results were then visualized as dendrograms and colored arrays in Java TreeView (University of Glasgow, UK).

### 2.4 Nervous cell type expression in mouse cortex

In order to assess the expression of the candidate genes on specific cell types of the nervous system, a search was made in the Brain RNA-Seq database (https://web.stanford.edu/group/barres_lab/brain_rnaseq.html), which is comprised of RNA sequencing data for acutely purified astrocytes, neurons, oligodendrocyte progenitor cells (OPC), newly formed oligodendrocytes (NFO), myelinating oligodendrocites (MO), microglia, and endothelial cells derived from the mouse cerebral cortex (Zhang et al., 2014). These data are available from our GitHub repository (https://github.com/lanec-unifesspa/gwas-pd/tree/master/RNA-Seq). Data were expressed as fragments per kilobase of exon per million fragments mapped (FKPM), and, after log transformation, were analyzed with hierarchical cluster with Euclidean distances and average linkage using Cluster 3.0 (University of Tokyo, Japan). Clustering results were then visualized as dendrograms and colored arrays in Java TreeView (University of Glasgow, UK).

### 2.5 Human-mouse disease connections

In order to evaluate cross-species disease connections of gene homologs, candidate genes were searched in the Human - Mouse: Disease Connection (HMDC) tool from the Mouse Genome Informatics database (Blake et al., 2017; http://www.informatics.jax.org). This tool aggregates data from mouse mutation, phenotype, and disease models with human gene-to-disease relationships from the National Center for Biotechnology Information (NCBI) and Online Mendelian Inheritance in Man (OMIM) and human disease-to-phenotype relationships from the Human Phenotype Ontology (HPO). Results were displayed as a matrix with all phenotypes/diseases associated with mouse models and human genes found for the candidate gene list.

### 2.6 Expression-phenotype correlations

For each gene discovered after filtering, an adequate probe within the well-curated INIA Amygdala Cohort Affy MoGene 1.0ST (Mar11) RMA, Hippocampus Consortium M430v2 (Jun06) PDNN, VCU BXD Prefrontal Cortex M430 2.0 (Dec06) RMA, INIA Hypothalamus Affy MoGene 1.0ST (Nov10), and INIA Adrenal Affy MoGene 1.0ST (Jun12) RMA Databases was identified using GeneNetwork (http://www.genenetwork.org; Williams and Mulligan, 2012)). These databases represent transcriptome datasets for different tissues of recombinant inbred mice. If several probes for the same gene were available, probes with higher maximum likelihood ratio statistic (LRS, a measurement of the association or linkage between differences in traits and differences in particular genotype markers values) were used.

For each gene, well-established threat conditioning paradigms, which represent good models for aspects of anticipatory anxiety and fear responses in panic attacks, were assessed in terms of correlation with expression in the aforementioned probes across recombinant lines. The following endpoints were included in the network analyses: Startle response to loud acoustic stimulus [force] (StartlePPI); Activity during first tone-shock pairing for males and females [units] (LM_PAIR1); Freezing response time following the final CS-US pairing during training in adult males and females [%] (LM_PAIRL); Freezing response time to conditioned cue after 24 hours (FearCued) [%]; Freezing response time to conditioned cue after 48 hours [%] (FearCue3); and Freezing response to context after 48 hours [%] (FearContext). As endocrinological trait, baseline control levels of corticosterone in plasma [ng/ml] (CORT BASE). Finally, both neuron (BLAmyNeurDensity) and glial cell (BLAmyGliaDensity) densities (n/mm^3^) in the basolateral amygdala (BLA) were selected as morphological traits. The association between expression levels from the GeneNetwork probes and these behavioral and morphological traits was analyzed with Pearson correlation coefficients. Results were represented graphically, and only traits showing Pearson correlation coefficients greater than 0.5 or less −0.5 were shown. These data are available from our GitHub repository (https://github.com/lanec-unifesspa/gwas-pd/tree/master/GeneNetwork).

### 2.7 Tests for enrichment of known indicated drugs

Drugs associated with each gene was extracted using the DsigDB database (Yoo et al., 2015; http://tanlab.ucdenver.edu/DSigDB/). The database was compiled according to bioassay results from PubChem and ChEMBL, kinase profiling assays from the literature and two kinase databases (Medical Research Council Kinase Inhibitor database and Harvard Medical School Library of Integrated Network-based Cellular Signatures database), differentially expressed genes after drug treatment derived from the Connectivity Map, and manually curated and text mined drug targets from the Therapeutics Targets Database and the Comparative Toxicogenomics Database. The resulting database holds gene sets for a total of 17839 unique compounds (as of February 2018). Each gene was searched in the database, and hit drugs were compiled in a spreadsheet. Following that, the Anatomical Therapeutic Classification (ATC) of each drug was search in the KEGG Drug database (https://www.genome.jp/kegg/drug/). Classification was annotated for the highest level (e.g., N [“Drugs acting on nervous system”] instead of N05 [“Psycholeptics”]). Enrichment tests were performed using Orange3 (Demsar et al., 2018, 2013), using the χ^2^ method to test for the association between ATC classification and gene.

## 3 Results

### 3.1 Metanalysis, prioritization, and candidate gene prediction

176 SNPs were found after the metanalysis. Visual inspection of Z scores suggested a bimodal distribution, with a peak around Z = −4.0 effect, and a second peak around Z = 4.25. Filtering SNPs based on FDR-corrected p-values yielded 25 SNPs significantly associated with PD. Z-scores for these filtered SNPs ranged from −5.177 to 4.6, with a median of −1.617 (Figure S1). PF-ranking generated a list of 29 genes with scores above 0 (Table S1). PF scores averaged 0.1051 (95% CI 0.0801 to 0.1301), similar to values found for candidate genes for breast cancer (Taşan et al., 2014). In GWASdb2 (Li et al., 2012), 134 genes with PF scores above 0 were found for the trait “panic disorder” (DOID: 594) (Table S2), averaging 0.0775 PF score (95% CI 0.0675 to 0.0875). The intersection between both datasets (Figure 1A) revealed 20 genes (*ACOT12*, *AHCY, ASB3, ASIP*, *CDK19*, *CENPI*, *CHAC2*, *CKMT2*, *CNTN4*, *EIF2S2*, *ERLEC1*, *GPR75*, *ITCH*, *PSME4*, *RASGRF2*, *SLC22A16*, *SOX21*, *TMEM35*, *TRMT2B*, and *ZCCHC9*) which were retained for further analyses. A moderate correlation between PF scores from the metanalysis and from GWASdb2 was found (r^2^ = 0.781; Figure 1B).

**Figure 1.**
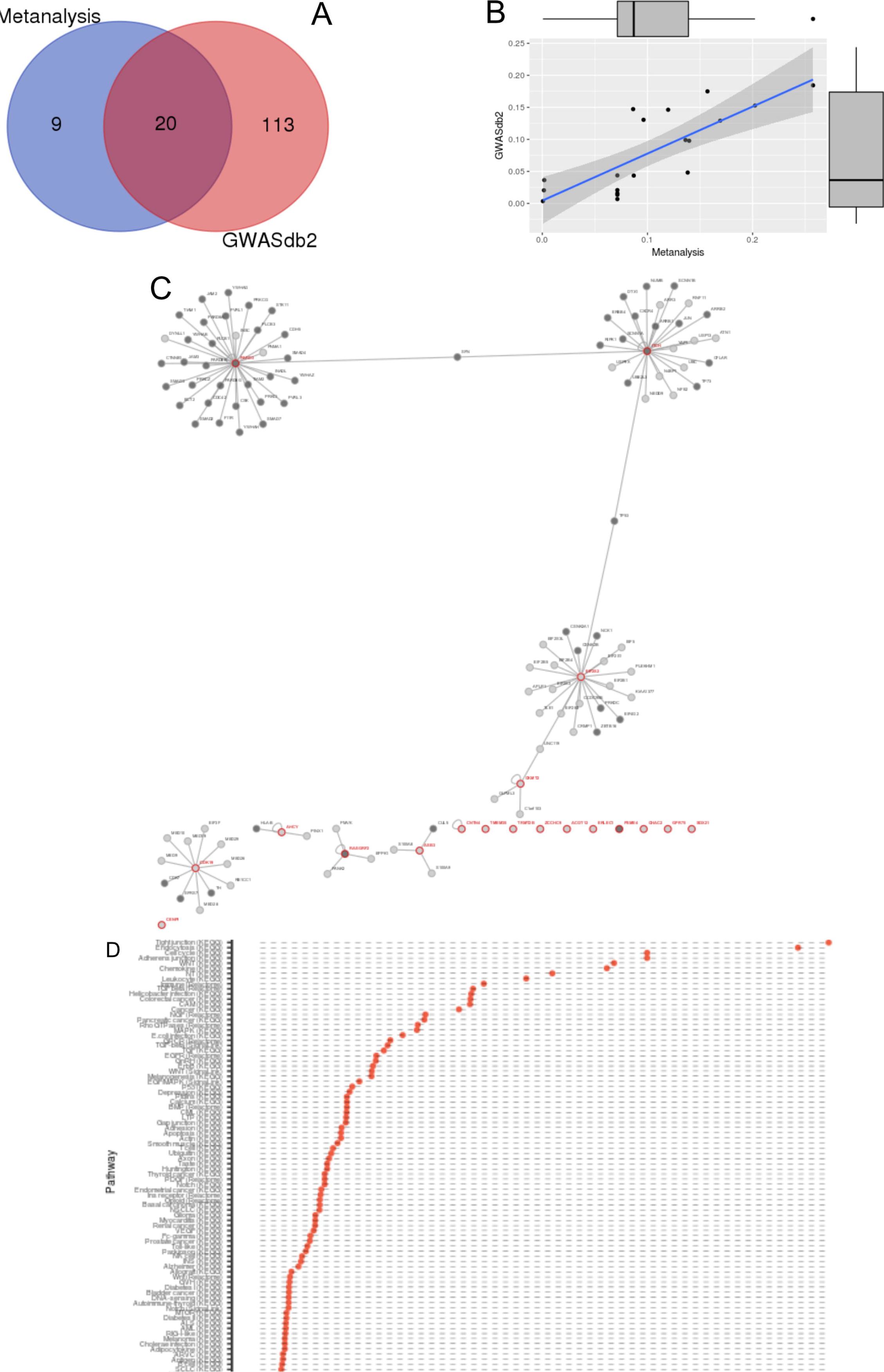
PrixFixe score ranking for prioritization of candidate genes in PD. (A) Venn diagram for candidate genes found after PF ranking in the metanalysis and candidate genes for PD predicted in GWASdb2. Genes in the intersection were kept for further analyses. (B) Correlation between PrixFixe scores found in the present study and in GWASdb2 (http://jjwanglab.org/gwasdb). The bands around the regression line represent 95% confidence intervals. The marginal boxplots represent values for PrixFixe scores for each dataset; the middle line indicates the medians, box limits indicate the 25th and 75th percentiles, and whiskers extend 1.5 times the interquartile range from the 25th and 75th percentiles. (C) Network of queried proteins (i.e., transcripts from the candidate genes) and their interactors. Red nodes represent the queried proteins, dark gray nodes represent signaling pathway member proteins, and light gray nodes represent non-pathway members. (D) Signaling pathways represented in the network. P-values below 0.001 are marked.

### 3.2 Protein-protein interaction and signalling pathway analyses

109 interactor proteins were found for these genes (Figure 1C; http://pathwaylinker.org/job.cgi?j=1511525998.96608266998). A major network (network N1) was formed by the interactors of *PARD3* (subnetwork S1), *ITCH* (subnetwork S2), *EIF2S2* (subnetwork S3), and *CKMT2* (subnetwork S4), linked by *SFN, TP53*, and *UNC119*. 3 other networks were found with more than 3 proteins: one involving *CDK19* (network N2), one involving *RASGRF2* (network N4), and one involving *ASB3* (network N5). *AHCY* participated in a small network, with 2 other proteins (network N3); 11 genes did not participate in any network.

Relevant signaling pathways in the interactor network included WNT (KEGG:hsa04310, Signalink), Chemokine (KEGG:hsa04062), Neurotrophin (KEGG:hsa04722), Immune system (Reactome:R-HSA-168256), TGF-beta (KEGG:hsa04350, Reactome:R-HSA-170834, Signalink), NGF (Reactome:R-HSA-166520), Rho GTPases (Reactome:R-HSA-194315), EGF/MAPK (KEGG:hsa04010, Signalink), Epidermal growth factor receptor (Reactome:R-HSA-177929), Gonadotropin-releasing hormone (KEGG:hsa04912), NGF (Reactome:R-HSA-166520), and ErbB (KEGG:hsa04012) (Figure 1D; http://pathwaylinker.org/job.cgi?j=1511525998.96608266998).

### 3.3 Brain and cell type expression

Expression patterns in the human brain were assessed via clustering of microarray data (Figure 2A). Clustering of brain regions revealed 3 clusters with |r^2^| ≥ 0.4. The first cluster (r^2^ = 0.648) included mainly limbic regions (frontal lobe, insula, cingulate gyrus, temporal lobe, striatum, claustrum, hippocampal formation, parahippocampal gyrus, amygdala, basal forebrain, epithalamus, and hypothalamus), as well as other cortical areas (occipital and parietal lobes). The second cluster (r^2^ = 0.716) included mainly motor regions (globus pallidus, ventral and dorsal thalami, subthalamus, mesencephalon, pontine tegmentum, myelencephalon, cerebellar nuclei, and basal part of the pons). The third cluster (r^2^ = 0.821) included white matter and sulci and spaces. Correlations for genes in the hierarchical cluster revealed 3 clusters with |r^2^| ≥ 0.4. The first cluster (r^2^ = 0.780), with low expression in the limbic cluster, moderate expression in the motor cluster, and very low expression in the white matter and sulci, included *ACOT12*, *ASIP*, *SLC22A16*, *CHAC2*, *TMEM35*, and *CNTN4*. The second cluster (r^2^ = 0.405) included *TRMT2B*, *ERLEC1*, *CENPI*, *CDK19*, *PSME4*, *EISF2S2*, and *ASB3*, low to moderate expression in the limbic cluster, low expression in the motor cluster, and moderate to high expression in the white matter and sulci. The third cluster (r^2^ = 0.527) included *CKMT2*, *ZCCHC9*, *ITCH*, *AHCY*, *SOX21*, and *GPR75*, which showed low expression in the limbic cluster, high expression in the motor cluster, and low expression in the white matter and sulci.

**Figure 2.**
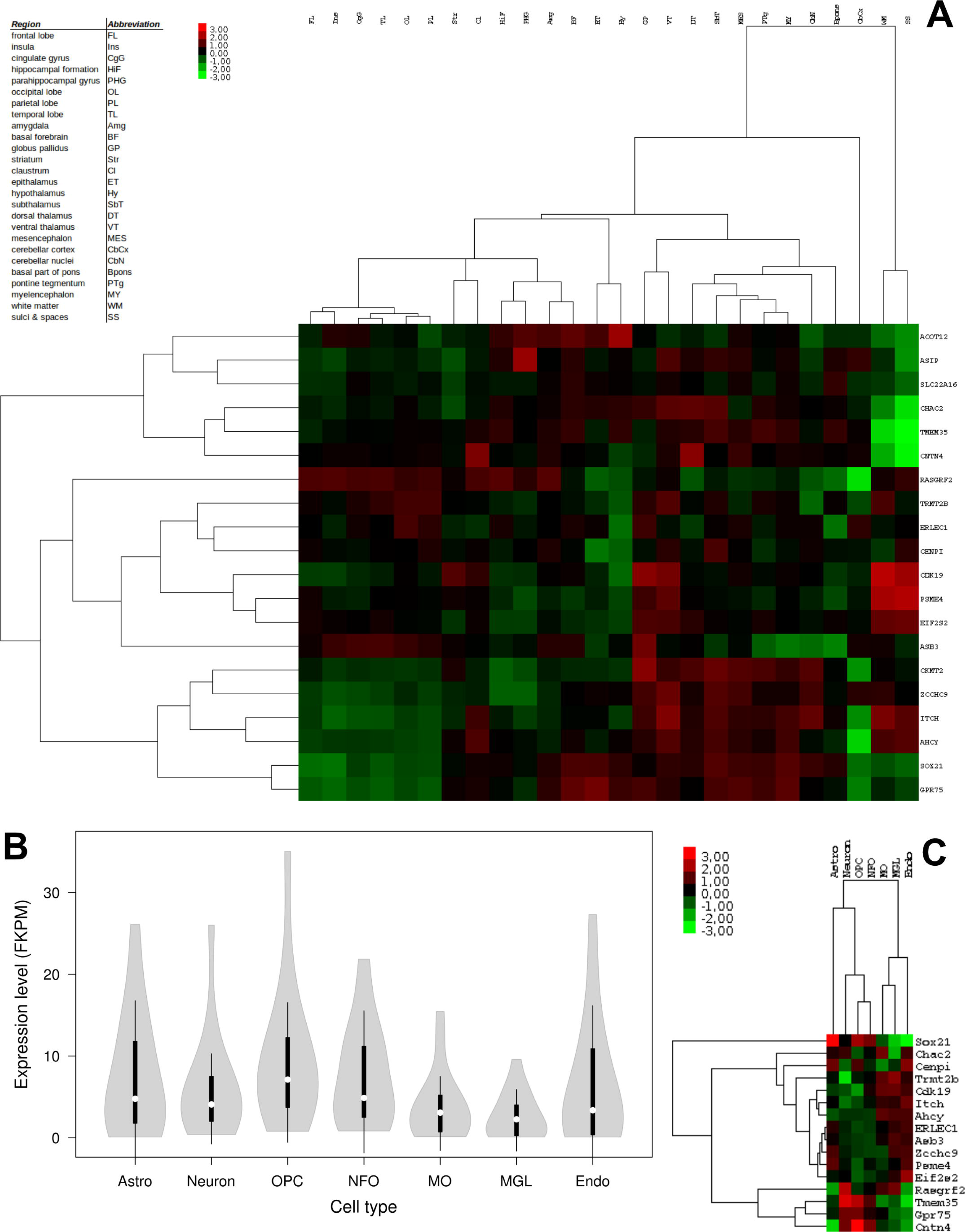
Expression patterns of candidate genes in the human brain (microarray data, A) and in mouse cortical cells (RNA-Seq data, B and C). The heatmap and cluster of human brain expression (A, http://human.brain-map.org/) was obtained by hierarchical clustering with Euclidean distances and average linking. (B) Violin plots representing the distribution of expression levels (in Fragments Per Kilobase Million [FKPM]) of the candidate genes in mouse cortical cells; white circles show the medians; box limits indicate the 25th and 75th percentiles; whiskers extend 1.5 times the interquartile range from the 25th and 75th percentiles; polygons represent density estimates of data and extend to extreme values. (C) Heatmap and cluster of RNA-Seq data in mouse cortical cells (https://web.stanford.edu/group/barres_lab/brain_rnaseq.html), obtained by hierarchical clustering with Euclidean distances and average linking.

16 of the 20 candidate genes were present in the Brain RNA-Seq database. In general, expression was higher in oligodendrocyte progenitor cells, newly formed cells, and astrocytes and neurons (Figure 2B). Hierarchical clustering suggested two clusters with |r^2^| ≥ 0.5 (Figure 2C). The first cluster (r^2^ = 0.676) included *Chac2*, *Cenpi*, *Trmt2b*, *Cdk19*, *Itch*, *Ahcy*, *Erlec1*, *Asb3*, *Zcchc9*, *Psme4*, and *Eif2e2*, with lower expression in astrocytes, neurons, OPCs, and NFOs, and higher expression in myelinating oligodendrocites (MOs), microglia, and endothelial cells. The second cluster (r^2^ = 0.556) included *Rasgrf2*, *Tmem35*, *Gpr75*, and *Cntn4*, with higher expression in astrocytes, neurons, OPCs, and NFOs, and lower expression in MOs, microglia, and endothelial cells.

### 3.4 Neuiobehavioial phenotypes in mice

Contrary to expectations, only two of the candidate genes were associated with behavioral or neurological abnormalities in mouse models (Figure 3). *Asip* was associated with abnormal food intake and hypoactivity; *Itch* was associated with neurological abnormalities (tremors and decreased grip strength). The lack of mouse models also demonstrate a literature gap, since only 12 of the 20 candidate genes were found in the MGI database. Moreover, it is probable that the published literature studying these genes in mouse mutants did not investigate behavioral phenotypes which are relevant to panic disorder.

**Figure 3.**
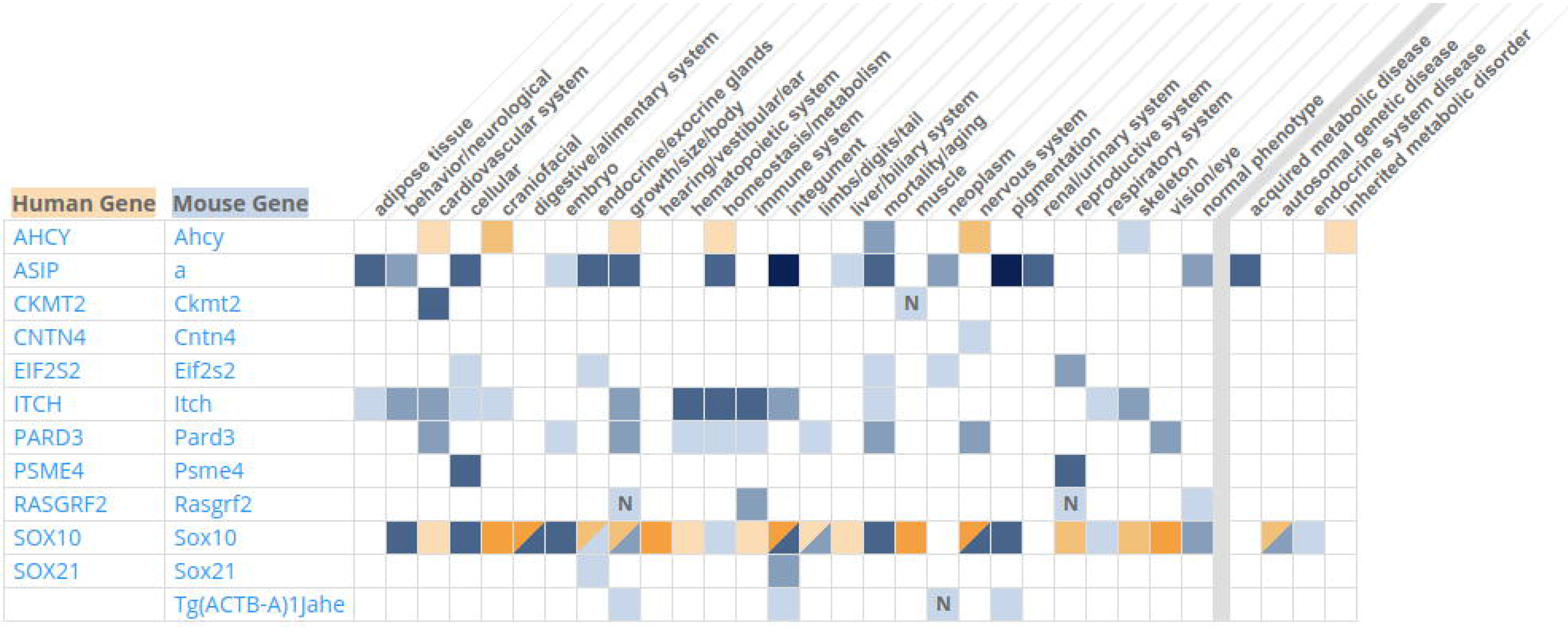
Human-mouse disease associations with candidate genes. Shading indicates term annotated to genes in humans (pink) or mice (blue); darker colors indicate more annotations. The letter “N” indicates lack of abnormalities in phenotype class in mice that contradicts expectations. Human gene-to-disease annotations are from NCBI and OMIM. Human phenotype-to-disease annotations are from HPO. Mouse gene-to-disease and gene-to-phenotype annotations are from MGI.

To further explore the associations between PD candidate genes and neurobehavioral phenotypes, we performed network analyses with mRNA expression data and behavioral traits in the BXD family of recombinant inbred strains. Behavioral and morphological traits showed moderate correlations between each in recombinant inbred mice lineages (Figures 4A-4E). BLA glial cell density was negatively correlated with baseline corticorsterone levels (r^2^ = −0.699) and positively correlated with startle response to loud acoustic stimulus (r^2^ = 0.523) and freezing response time to conditioned cue after 24 (r^2^ = 0.560) and 48 h (r^2^ = 0.685). BLA neuron cell density was negatively correlated with freezing response time to context after 48 h (−0.607) and with baseline corticosterone levels (r^2^ = −0.657). Freezing response time following the final CS-US pairing during training was positively correlated with freezing to conditioned cue after 24 h (r^2^ = 0.543).

**Figure 4.**
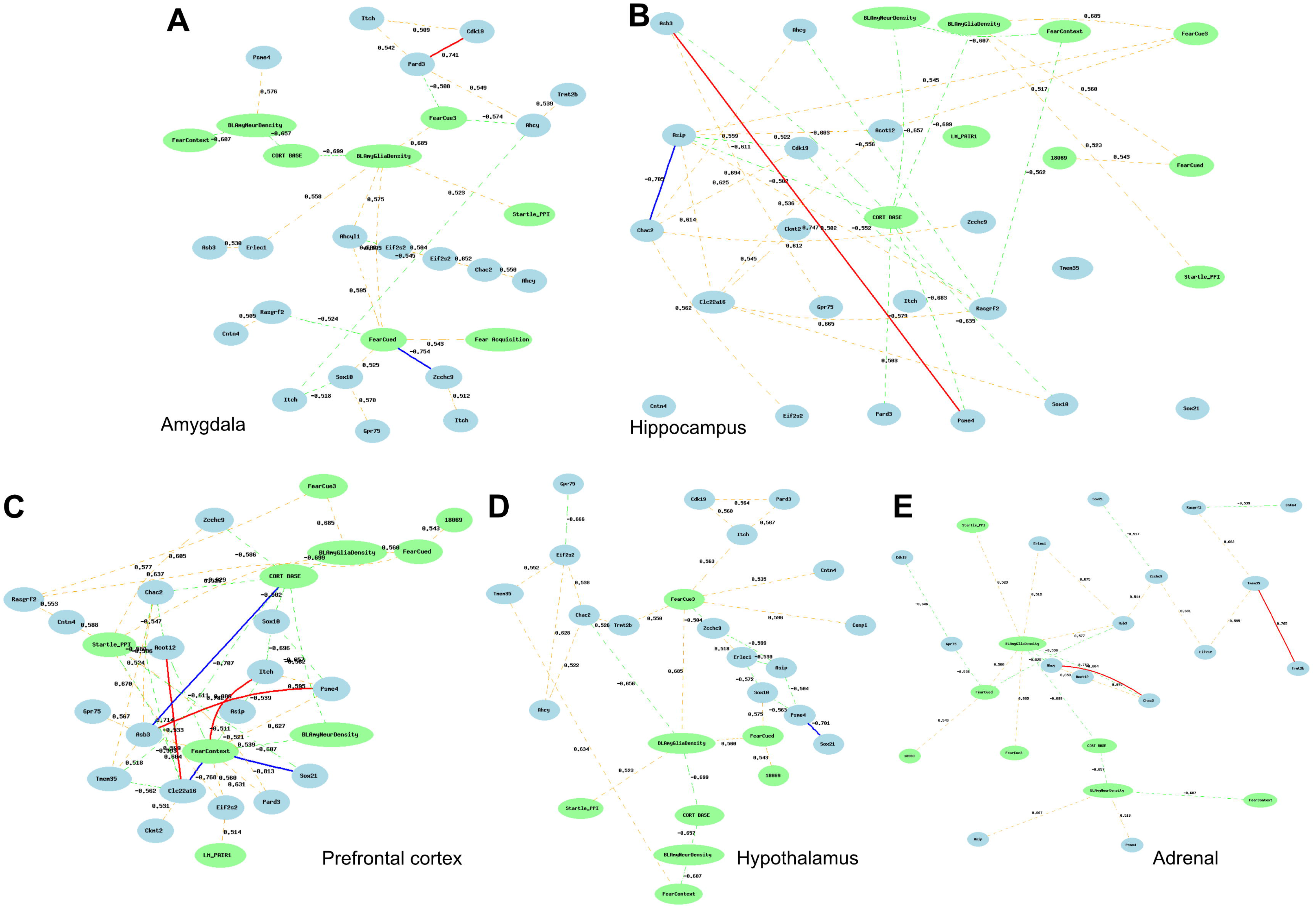
Correlations between mRNA expression in (A) amygdala, (B) hippocampus, (C) prefrontal cortex, (D) hypothalamus, and (E) adrenal gland and behavioral, endocrinological, and morphological traits in BXD mice. Blue ellipses represent genes in the dataset, and green ellipses represent the neurobehavioral traits. Full lines connecting edges represent |r^2^| ≥ 0.7 (red = positive, blue = negative), and dashed lines represent 0.5 ≤ |r^2^| ≤ 0.7 (orange = positive; green = negative).

In the amygdala mRNA probe set, a network with 20 nodes showing |r^2^| ≥ 0.5 was formed (Figure 4A). A strong negative correlation between *Zcchc9* mRNA levels in the amygdala across inbred lineages and freezing to conditioned cue after 24 h (r^2^ = −0.754) was found. Weaker correlations were found between amygdalar *Erlec1* mRNA levels and BLA glial density (r^2^ = 0.505); between amygdalar *Psme4* mRNA levels and BLA neuron density (r^2^ = 0.576); and between freezing to conditioned cue after 24 h and amygdalar *Rasgrf2* (r^2^ = 0.539).

In the hippocampus mRNA probe set, a network with 25 nodes showing |r^2^| ≥ 0.5 was formed (Figure 4B), with moderate negative correlations between basal corticosterone levels and hippocampal *Pard3* (r^2^ = −0.579), *Psme4* (r^2^ = −0.684), *Asip* (r^2^ = −0.502), and *Asb3* (r^2^ = −0.603) mRNA levels; moderate positive correlations between freezing to conditioned cue after 48 h and hippocampal *Acot12* (r^2^ = 0.517) and *Asip* (r^2^ = 0.545) mRNA levels; and a moderate negative correlation between freezing to context after 48 h and *Rasgf2* mRNA levels in the hippocampus across lines (r^2^ = −0.562).

In the prefrontal cortex mRNA probe set, a network with 25 nodes showing |r^2^| ≥ 0.5 was formed (Figure 4C), with a strong negative correlation between basal corticosterone levels and *Asb3* mRNA levels in the PFC (r^2^ = −0.707); strong positive correlation between freezing to context after 48 h and PFC *Itch* mRNA levels (r^2^ = 0.806), and negative correlations with PFC *Slc22a16*(r^2^ = −0.768) and *Sox21* (r^2^ = |0.813) levels; moderate positive correlations between startle response to loud acoustic stimulus and PFC *Cntn4* (r^2^ = 0.588), *Zcch9* (r^2^ = 0.605), and *Eisf2s2* (r^2^ = 0.678) mRNA levels, and negative with *Slc22a16* (r^2^ = −0.533) levels in the PFC; a moderate positive correlation between freezing to conditioned cue after 24 h and PFC *Rasgrf2* mRNA levels (r^2^ = 0.637); and a moderate positive correlation between freezing to conditioned cue after 48 h and PFC *Rasgrf2* mRNA levels (r^2^ = 0.605).

In the hypothalamus mRNA probe set, a network with 25 nodes showing |r^2^| ≥ 0.5 was formed (Figure 4D). A moderate negative correlation was found between BLA glial cell density and hypothalamic *Chac2* mRNA levels (r^2^ = −0.656). Moderate positive correlations was found between freezing to conditioned cue after 48 h and hypothalamic *Trmt2b* (r^2^ = 0.550), *Itch* (r^2^ = 0.563), and *Cntn4* (r^2^ = 0.535) mRNA levels. A moderate positive correlation was found between freezing to context and *Tmem35* mRNA levels in the hypothalamus (r^2^ = 0.634).

In the adrenal gland mRNA probe set, a network with 24 nodes showing |r^2^| ≥ 0.5 was formed (Figure 4E). Moderate positive correlations were found between BLA neuron density and adrenal *Asip* (r^2^ = 0.667) and *Psme4* (r^2^ = 0.518) mRNA levels, and between BLA glial cell density and adrenal *Erlec1* (r^2^ = 0.512) mRNA levels. Negative moderate correlations were found between freezing to conditioned cue after 24 h and adrenal *Gpr75* (r^2^ = −0.558) and *Asb3* (r^2^ = −0.536) mRNA levels.

### 3.5 Drug repositioning

Top 20 drugs hits after searching the DSigDB database can be found in Figure 5A. No ATC classes were enriched in the set (χ^2^ = 740.62, p = 1, N = 279). No significant enrichment was observed at the drug level (χ^2^ = 5800.41, p = 0.146, N = 549)(Figure 5B).

**Figure 5.**
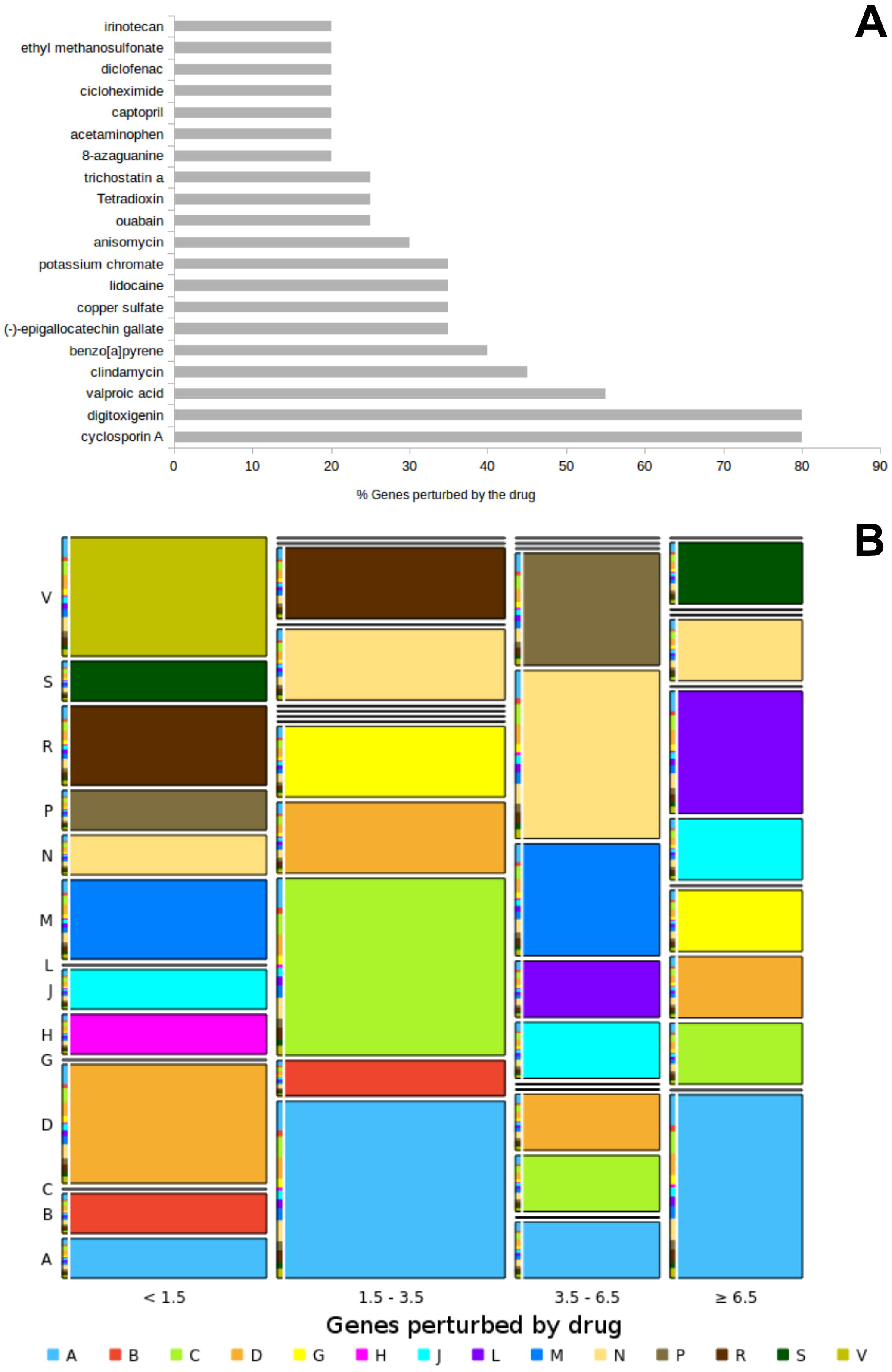
(A) Top 20 hits on DSigDB for causal genes for PD. (B) Mosaic plot of drug classes (Anatomical Terminology Classification) found in the set, separated by gene/protein that is perturbed with treatment. Letters are as follows: A – Alimentary tract and metabolism; B – Blood and blood forming organs; C – Cardiovascular system; D – Dermatologicals; G – Genito-urinary system and sex hormones; H – Systemic hormonal preparations; J – Antiinfectives for systemic use; L – Antineoplastic and immunomodulating agents; M – Musculo-skeletal system; N – Nervous system; P – Antiparasitic products, insecticides and repellents; R – Respiratory system; S – Sensory organs; V – Various.

## 4 Discussion

### 4.1 PF ranking

None of the candidate genes found here figured in a dataset for six major neuropsychiatric disorders (attention deficit hyperactivity disorder, anxiety disorders, autistic spectrum disorders, bipolar disorder, major depressive disorder, and schizophrenia) (Lotan et al., 2014). Notably, none of the candidate genes found here figured in a dataset of candidate genes (derived from both human and murine GWAS) for anxiety-spectrum phenotypes, including anxiety disorders, neuroticism, and major depression, as well as anxiety-like behavior in mice (Hettema et al., 2011).

### 4.2 Protein-protein interactions and functional associations

When mapping the protein-coding candidate genes to a merged and curated BioGRID and HPRD protein-protein interaction database, only 109 proteins, with no interactions between them, were found. Similarly to a cross-disorder set of candidate genes (including genes for ADHD, autistic spectrum disorders, bipolar disorder, major depressive disorder, and schizophrenia; Lotan et al., 2014), thus, the PD dataset does not form an interconnected network at the level of direct protein-protein interactions. Despite the lack of direct interactions, core proteins from each network were connected via second interactors for three of the 20 protein-coding genes of the set (ITCH, EIF2S2, and CKMT2). These interactor proteins were involved in the interaction of multiple signaling pathways (SFN), signal transduction in immune cells (UNC119), and cell cyle (TP53). These links were not surprising, since these molecules are overrepresented in the genetics literature (Dolgin, 2017), and since they participate in a wide array of interactions (SFN: n = 126; TP53: n = 342; UNC119: n = 84); moreover, these proteins have not yet been associated with neuropsychiatric disorders. Therefore, interaction is unlikely to be specific for PD.

In spite of this lack of direct protein-protein interaction, important signaling pathways were found to be enriched in the PD dataset and interactor network – including WNT, immune and inflammatory pathways, neurotrophins, EGF/MAFPK, and Rho GTPase pathways. Nonetheless, from the candidate genes only *RASGRF2* (which was part of the MAPK, NGF, and Rho GTPase pathways) and *ITCH* (which was part of Endocytosis, Ubiquitin, and NGF pathways) were part of enriched signaling pathways. Both the MAPK and the Wnt pathways were implicated in threat conditioning and aversive learning in animal models (Maguschak and Ressler, 2011; Schafe et al., 2001). NGF, on the other hand, has been implicated in stress and stress-related psychopathology (Cirulli and Alleva, 2009). While most of these pathways have been implicated in neural development, mRNA expression of PF-ranked genes in recombinant inbred mice was poorly associated with morphological traits of the BLA; moreover, these associations were more common between BLA morphology and mRNA expression of candidate genes in the HPA axis, and *not* in the amygdala, as would be expected. Interestingly, *Erlec1* and *Psme4* expression in both the amygdala and adrenal were associated with BLA morphology.

### 4.3 Tissue and cell expression

Expression patterns in the human brain suggested that most genes were at best moderately expressed in “classical” limbic regions (frontal and cingulate cortices, hippocampus, amygdala); higher expression was found for *TRMT2B*, *ERLEC1*, *CENPI*, *CDK19*, *PSME4*, *EISF2S2*, and *ASB3*. In rodent PFC cell types, orthologues *Trmt2b*, *Eisf2s2*, *Cdk19*, *Erlec1*, and *Asb3* show higher expression in MOs, microglia, and endothelial cells. These results contrast with a recent GWAS of major depression, in which genes were highly expressed in the frontal and cingulate cortices, hippocampus, and basal ganglia, and significantly enriched in neurons (vs. oligodendrocytes and astrocytes) (Wray et al., 2018), suggesting disorder-specific patterns. While a region-specific pattern of expression cannot be discarded, these results suggest that MOs and microglial cells in limbic regions are associated with PD. Among these genes only CDK19 and ASB3 appeared in interactor networks (CDK19 was the core of N2), and none was part of the enriched signaling pathways; overall, these results suggest that expression patterns, instead of general function in interactor networks and signalling pathways, is the commonality between these PD-associated genes.

When mRNA expression across limbic regions is associated with behavioral traits in rodents, a more specific pattern emerges. Some genes were associated with behavioral traits in fear conditioning paradigms; in all cases, the association was with freezing to cue and/or to context. Gene expression in the amygdala (*Zcchc9*, *Rasgrf2*, *Sox10*) was associated with freezing to cue 24 h after conditioning, while gene expression in the hypothalamus (*Trmt2b*, *Itch*, *Cntn4*) was associated with freezing to cue 48 h after conditioning. Genes expression in the PFC (*Itch*, *Slc22a16*, *Sox21*) were negatively associated with freezing to context 48 h after conditioning. In the mouse PFC, *Zcchc9*, *Itch*, and *Trmt2b* were expressed mainly in MOs, microglia, and endothelial cells, while *Rasgrf2* and *Cntn4* were expressed mainly in astrocytes, neurons, OPCs, and NOFs.

The higher expression in MOs is of interest, since myelin is known to stabilize synaptic networks, and oligodendrocyte-related proteins have been shown to be modified in the rat dentate gyrus during contextual fear conditioning (Houyoux et al., 2017). Moreover, microglia has been shown to participate in cued fear conditioning (Parkhurst et al., 2013), and microglial acid sensing has been shown to participate in carbon dioxide-evoked fear (Vollmer et al., 2016). These results also make sense in light of the enrichment of immune and inflammatory pathways in the interactor network.

### 4.4 Drug repositioning attempts

Top 20 drugs which perturb PF-ranked genes include interesting findings. Cyclosporin A has been shown to inhibit calcineurin (Fakata et al., 1998), impairing aversive memory formation in chicks (Bennett et al., 1996); and valproic acid enhances the extinction and habituation of fear conditioning in healthy humans (Kuriyama et al., 2011). Nonetheless, evidence for their use in treating PD (or in exacerbating PD symptoms) is scarce. The current results do not provide sufficient evidence for the association of these drugs and their effects on the PF-ranked genes found for PD; in fact, no gene-drug association was found either at the individual drug level, or at the drug class level. Differently from previous reports (So, 2017; So et al., 2016), which support drug repositioning from genes extracted from GWAS for anxiety and mood disorders, our findings do not support such proposals. Extensive methodological differences are probably responsible for this discrepancy.

### 4.5 Consequences for the pathophysiology and pharmacology of PD

One important limitation of the present results is that they are exploratory in nature, awaiting confirmatory research on all findings. Nonetheless, the present results suggest interesting avenues of investigation on the relationship between genetics and the pathophysiology and pharmacology of PD. First, the observation of cross-species associations between genes discovered in the GWAS metanalysis and neurobehavioral traits of fear prompts the idea of conservation across species of specific endophenotypes (de Mooij-van Malsen et al., 2011; Kas et al., 2012), such as alterations in fear conditioning (Battaglia and Ogliari, 2005; Bouton et al., 2001). Second, and perhaps a more interesting point, is the observation that these endophenotypes are not only related to the expression levels of risk genes in regions such as prefrontal cortex, amygdala, and hypothalamus, but also that these genes are more expressed in myelinating oligodendrocytes and microglia than neurons and astrocytes, the brain cells which are more studied and usually highly associated with psychiatric disorders in GWAS ontology studies (Lotan et al., 2014; Wray et al., 2018).

This last observation ties well with the observation of enrichment in immune and inflammatory pathways, given the role of microglia in these functions in the brain (Hanisch, 2002). Small studies have suggested increased levels of proinflammatory cytokines in serum from patients with PD (e. g., Hoge et al., 2009), and cytokines have been associated with acute mental stress in non-clinical populations (Kunz-Ebrecht et al., 2003; Porterfield et al., 2011; Steptoe et al., 2001). Whether these changes are generalized in PD, and whether genes associated with increased PD risk do so by altering peripheral cytokine responses as well as microglia and oligodendrocyte function in the brain is still unknown.

The present results also highlight putative targets for basic pharmacological research. While drug repositioning analysis was inconclusive, experimental drugs acting on these pathways were not part of the dataset. Therefore, the present results point towards an interesting *a posteriori* hypothesis that targeting specific pathways (WNT, immune and inflammatory responses, neurotrophins, EGF/MAPK, Rho GTPase) specifically in microglia and oligodendrocytes could elucidate the pathophysiology of PD and suggest novel investigational drugs. Further confirmatory research will allow that hypothesis to be tested.

---

Figure S1 – (Upper panel) Distribution of unfiltered z-values of SNPs found in the metanalysis. (Lower panel) Distribution of z-values for the metanalysis after filtering for the false discovery rate.

